# TreeMap: A Structured Approach to Fine Mapping of eQTL Variants

**DOI:** 10.1101/2020.05.31.125880

**Authors:** Li Liu, Pramod Chandrashekar, Biao Zeng, Maxwell D. Sanderford, Sudhir Kumar, Greg Gibson

## Abstract

**Motivation:** Expression quantitative trait loci (eQTL) harbor genetic variants modulating gene transcription. Fine mapping of regulatory variants at these loci is a daunting task due to the juxtaposition of causal and linked variants at a locus as well as the likelihood of interactions among multiple variants. This problem is exacerbated in genes with multiple cis-acting eQTL, where superimposed effects of adjacent loci further distort the association signals.

**Results:** We developed a novel algorithm, TreeMap, that identifies putative causal variants in cis-eQTL accounting for multisite effects and genetic linkage at a locus. Guided by the hierarchical structure of linkage disequilibrium, TreeMap performs an organized search for individual and multiple causal variants. Via extensive simulations, we show that TreeMap detects co-regulating variants more accurately than current methods. Furthermore, its high computational efficiency enables genome-wide analysis of long-range eQTL. We applied TreeMap to GTEx data of brain hippocampus samples and transverse colon samples to search for eQTL in gene bodies and in 4 Mbps gene-flanking regions, discovering numerous distal eQTL. Furthermore, we found concordant distal eQTL that were present in both brain and colon samples, implying long-range regulation of gene expression.

**Availability:** TreeMap is available as an R package enabled for parallel processing at https://github.com/liliulab/treemap.

## INTRODUCTION

Scans for expression quantitative trait loci (eQTL) aim to discover genetic variants associated with variation in transcript abundance among individuals. Genome-wide scanning of eQTL involves genomic and transcriptomic profiling of a large number of samples, followed by statistical and experimental analyses of polymorphic sites to discover expression quantitative trait nucleotides (eQTNs) (1). Due to linkage disequilibrium (LD), typically multiple genetic variants at a locus show highly significant statistical scores, although only some of these are causal eQTNs. Expression-associated variants (eVars) are usually aggregated into a credible set that includes a lead variant with the strongest association signal and other linked variants. However, a lead eVar is not necessarily responsible for transcriptional regulation, but tags causal eQTNs instead (2,3). Furthermore, in genes with multiple cis-acting eQTL, the correspondence between lead eVars and causal variants diminishes quickly due to superimposed effects of adjacent loci (4–6).

To better resolve causal variants, recent fine-mapping efforts have gone beyond the conventional single-locus assumption and evaluated multi-locus effects (7–10). Because an exhaustive search for an unknown number of causal variants in a wide genomic region is computationally prohibitive, several strategies have been employed to ease the computational burden. Stepwise conditional regression is a greedy algorithm that repeatedly tests individual sites and returns lead eVARs with the best marginal test statistics at each iteration (10). This algorithm is computationally efficient although the solution is highly susceptible to local optima. To overcome local optima, the piMASS method performs a Markov chain Monte Carlo search for a user-specified number of causal variants (11). However, the requirement of prior knowledge of the number of causal variants and the high computational cost make it impractical for genome-wide analysis. The adaptive DAP method takes a tiered strategy that first scans a genomic region for independent eQTL and then conducts an exhaustive search within each locus (7). Although this method does not impose constraints on the number of causal variants, attempts at finding more than four causal variants are still computationally intensive (5). Given that most human genes have multiple cis-acting eQTL (12,13) and independent studies have reported that credible intervals generally contain one hundred or more eVars per gene (14–16), fine-mapping algorithms capable of identifying an arbitrary number of eQTL, prioritizing multiple eVars at a locus, and performing at high computational efficiency will improve genome-wide discovery of regulatory variants.

While LD between eVars adds to the complexity of eQTL fine-mapping, it also provides a convenient structure with which large genome regions can be dissected into multiple relatively independent segments that are then amenable to association testing. The Tree Scanning method uses an evolutionary tree of haplotypes to study phenotypic associations (17). However, because this method uses haplotypes as the genomic unit, it lacks base-pair resolution and is unsuited to fine-mapping tasks. Tree-guided lasso (18) offers an intuitive solution, in which selection of groups of variants or individual variants is conducted in a hierarchical framework defined by LD structure. This machine-learning method is also highly efficient for genome-scale analysis. However, it does not provide statistical confidence on the selected features required for biological and clinical applications.

To address these deficiencies, we designed a nested model that first employs the tree-guided lasso algorithm to scan a large genomic region for candidate loci and candidate variants within a locus, and then apply statistical inference to derive credible sets of putative causal variants. We tested this new method, named TreeMap, via rigorous simulations. We show that TreeMap has significantly higher accuracy and faster computation than existing methods under various scenarios, especially for genes with multiple cis-acting eQTL under weak to medium LD. Applications of TreeMap to GTEx data of brain hippocampus samples and transverse colon samples revealed abundant distal regulatory variants located in up to 2 Mbps away from gene bodies.

## MATERIALS AND METHODS

### Data structure

Given *n* samples, each genotyped at *m* biallelic positions in the upstream region of a target gene, a feature matrix *X* contains genotype data with rows corresponding to samples and columns corresponding to genetic variants. A response vector *Y* contains expression level of the target gene in *n* samples. To represent the LD structure of the variants, we compute the squared correlation coefficient (*r*^2^) between pairs of variants. We define six *r*^2^ cutoffs (>0.999, 0.98, 0.95, 0.90 0.85 and 0.80). Using each cutoff, we convert the correlation matrix into an adjacency matrix and construct an undirected graph with the greedy clustering algorithm (19). During the clustering process, we reserve the order of neighboring variants and require the largest within-cluster gap <100 consecutive variants. Each cluster in the graph represents an LD block for a specific *r*^2^ cutoff. We then organize these blocks into a hierarchical structure *G* with 8 levels (**Fig. 1A**). At the leaf level (*G*^0^), each node represents a single variant. At higher levels in a sequential order (*G*^1^, …, *G*^6^), each node represents variants belonging to an LD block with *r*^2^ >0.999, 0.98, 0.95, 0.90 0.85 and 0.80, respectively. The root level (*G*^7^) has a single node containing all variants.

**Figure 1.**
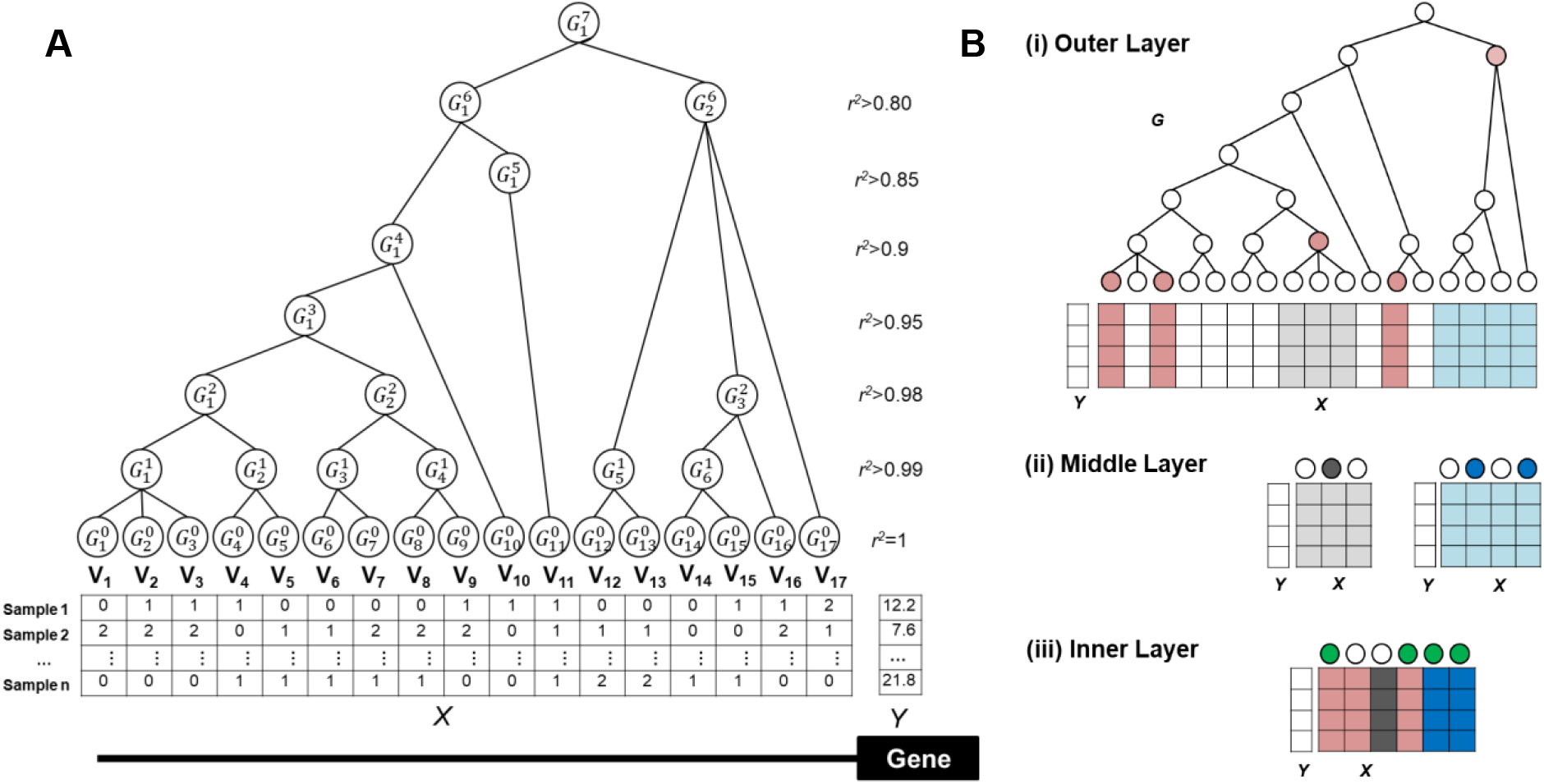
The TreeMap method. (**A**) Data structure. *X* is a feature matrix containing genotypes of *V* variants. *Y* is a response vector containing transcriptional abundances of the target gene. Variants are organized into a hierarchical structure *G* that reflects different levels of linkage estimated by *r^2^* values. (**B**) Nested design. (**i**) At the outer layer, individual variants (leaf nodes, red circles) or groups of variants (internal nodes, red circles) associated with gene transcription are selected. (**ii**) At the middle layer, variants belonging to the selected groups (gray blocks and blue blocks) are tested for node-specific optimal solutions (dark gray circles and dark blue circles). (**iii**) At the inner layer, variants selected from previous layers are aggregated to identify a global optimal solution (green circles).

### TreeMap framework

TreeMap takes a 3-layer nested design to remove uninformative variants and reduce redundancies among informative variants progressively (**Fig. 1B**). At the outer layer, tree-guided penalized regression selects groups of variants (internal nodes in *G*) or individual variants (i.e., leaf nodes in *G*) associated with transcriptional changes. At the middle layer, stepwise conditional multivariate tests iterate combinations of variants within each selected node to identify a node-specific optimal solution. At the inner layer, variants selected from the previous layers are aggregated and passed through a Bayesian multivariate analysis to derive a global optimal solution. The final solution satisfies both between-locus sparsity by selecting only a few internal nodes, and within-locus sparsity by selecting only a few individual variants in a node. Below we provide detailed descriptions of each layer.

#### Outer layer

We formulate the selection of causal variants from a genomic region with LD structure as a sparse learning problem under graph constraints. Specifically, given a feature matrix *X* with *n* rows and *m* columns, a response vector *Y* of length *n*, and a hierarchical relationship *G* of features in *X* with *d* levels, we will learn a linear model *Y* = *Xβ* + *ϵ* that solves

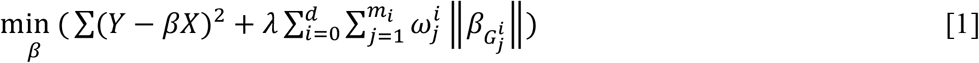

where *β* is a vector of coefficients of individual variants, 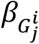 is the vector of coefficients of variants belonging to a node 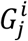, *λ* is the regularization parameter, and 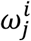 is the weight of each node in group 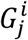. We compute 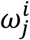 as

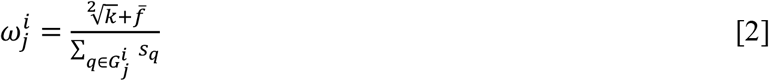

where *k* is the number of variants in the group, 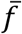 is the average minor allele frequency, and *s_q_* is a user-specified functional impact score of a variant *q* in the group.

The sparsity (i.e., the number of variants with non-zero *β* values) of the solution to equation [1] is controlled by *λ*, such that a larger *λ* value leads to fewer selected variants. In practice, choosing the most appropriate value of *λ* is mostly subjective. To address this problem, we test a range of *λ* values with bootstrap samples and regard the top 5% variants receiving non-zero *β* values as informative. The *β* value of an internal node is the average of its member variants. The top 5% internal nodes receiving non-zero *β* values are also informative. We denote the set of variants selected at this layer as *S*_1_.

#### Middle layer

For each informative internal node, we perform a stepwise conditional analysis to find a set of variants in *S*_1_ with non-redundant information. Specifically, given a node containing a set of variants *V*, we first fit a linear regression model for each and every member variant *q* as

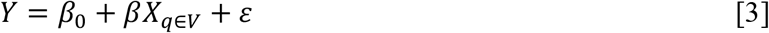

Among all member variants passing a statistical threshold (i.e. Bonferonni-corrected p value <0.05 and explained residual >1%), we choose the variant with the smallest p-value as the primary variant. Next, conditional on this primary signal, we test each remaining variant by fitting a linear regression model on the residual *ε* and identify the variant with the smallest p-value. We repeat this process until exhausting all member variants or no remaining variants passing the statistical threshold. We then aggregate variants selected from this procedure with variants in *S*_1_ that do not belong to any informative internal nodes, and map them into nodes at the *G*^6^ level (i.e., *r*^2^ > 0.8). Within each node, we iterate all combinations of one or two variants to fit a linear regression model and compute the Akaike information criterion (AIC) values

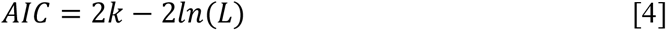

where *k* in the number of variants included and *L* is the likelihood of the fitted model. We select the variants giving rise to the smallest AIC value and denote this set as *S*_2_.

#### Inner layer

If *S*_2_ contains no more than 10 candidate variants, we perform an exhaustive search for the best linear model with an arbitrary number of variants based on the Bayes factor. We define *M* as a multivariate linear model with selected variables and *M*_0_ as a null model with no independent variables. By giving equal prior probabilities to *M* and *M*_0_, the *BF* is

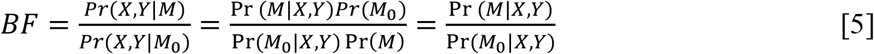

The set of variants giving rise to the largest *BF* value constitutes the lead variants of the credible set. If *S*_2_ contains more than 10 candidate variants, we use backward stepwise selection based on AIC values as in equation [4] to identify lead variants. Using each lead variant as an anchor, we scan *S*_1_ for tagging variants with *r*^2^>0.5 linked to the lead variant. We define an eQTL as a lead variant with its tagging variants ranked on *r*^2^ values. The final credible set may contain multiple loci.

#### Estimate effect sizes

After we derive a final credible set for a gene, we build a linear regression model

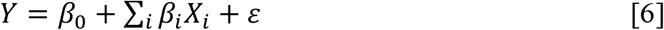

where *Y* is the transcript abundance, *X_i_* is the lead eVar of the *i^th^* eQTL and *β_s_* are effect sizes, and *ε* is errors. For each *X_i_*, we test the null hypothesis of *β_i_* = 0 and use the p-value to represent the statistical significance of the corresponding eQTL. We consider the eQTL with the best p-value as the primary locus and the remaining eQTL as auxiliary loci.

### Simulation data

We used an established approach (5) to simulating gene transcription controlled by one to ten causal variants. Given a randomly picked human gene, we retrieved genotypes *X* of all variants located in the 200 kb upstream region of its transcription start site from the 1000 Genomes Project phase 3 data (20). From among these variants, we picked *h* random variants as causal variants, and assigned each causal variant *i* an effect size 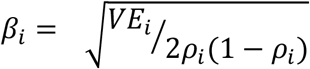 where *ρ_i_* is the minor allele frequency, and *VE_i_* is the variance explained. We allowed *VE_i_* to take a random value from a uniform distribution *unif*(0.02, 1). We then simulated gene transcript abundance 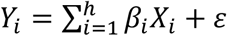 where *ε* is the environmental noise following a normal distribution *norm*(0, 1). On average, each simulation involved 1,700 variants genotyped in 1,835 samples with non-African ancestry from the 1000 Genome Project.

### GTEx datasets

We downloaded RNA-seq data of 123 brain hippocampus samples and 274 transverse colon samples from the GTEx data portal (v7, mapped to the hg19 reference genome) (21). Transcript abundance quantified as Transcripts Per Kilobase per Million mapped reads (TPM) were available for 23,725 genes in brain and for 24,423 genes in liver. Following the recommendations from the GTEx Consortium, we adjusted TPM values for technical covariates using multivariate linear regression (22). For each gene, we retrieved genotypes of common variants (minor allele frequency MAF > 0.05) located inside the region from 2 Mbps upstream of the transcription start site (TSS) to 2 Mbps downstream of the transcript end site (TES). On average, each gene had 8,199 common variants. We built a hierarchical tree of these variants using the method described above and applied TreeMap to each gene.

## RESULTS

Using simulation data, we tested the performance of TreeMap, DAP and stepwise conditional analysis. We then applied TreeMap to analyze GTEx data of brain hippocampus samples and transverse colon samples.

### Performance on computer simulations

#### Mapping independent causal variants

We randomly sampled 400 genes from the human genome and simulated 1, 2, 3, and 4 causal variants for each gene. We required that r^2^ values between all pairs of causal variants of a gene were less than 0.1. These simulations represented genes with only independent cis-acting eQTL.

We first examined if each method reported the correct number of independent eQTL. When a gene had a single causal locus, TreeMap reported the correct number 98% of the time, which was significantly higher than DAP (94%, two-proportion test *P*=0.003) and conditional analysis (75%, *P*=10^−21^). As the number of independent causal loci per gene increased, the accuracies of all three methods decreased linearly (**Fig. 2A**). However, the accuracy of TreeMap remained as the highest among the three methods in all scenarios. For genes with four independent causal loci, TreeMap still made correct predictions 79% of the time, whereas the accuracies of DAP and conditional analysis dropped to 70% (*P*=0.002) and 58% (*P*=10^−9^), respectively. When these methods made wrong predictions, they tended to over-estimate the number of independent causal loci, with conditional analysis showing the largest deviations and TreeMap showing the smallest deviations (**Fig. 2B**).

**Figure 2.**
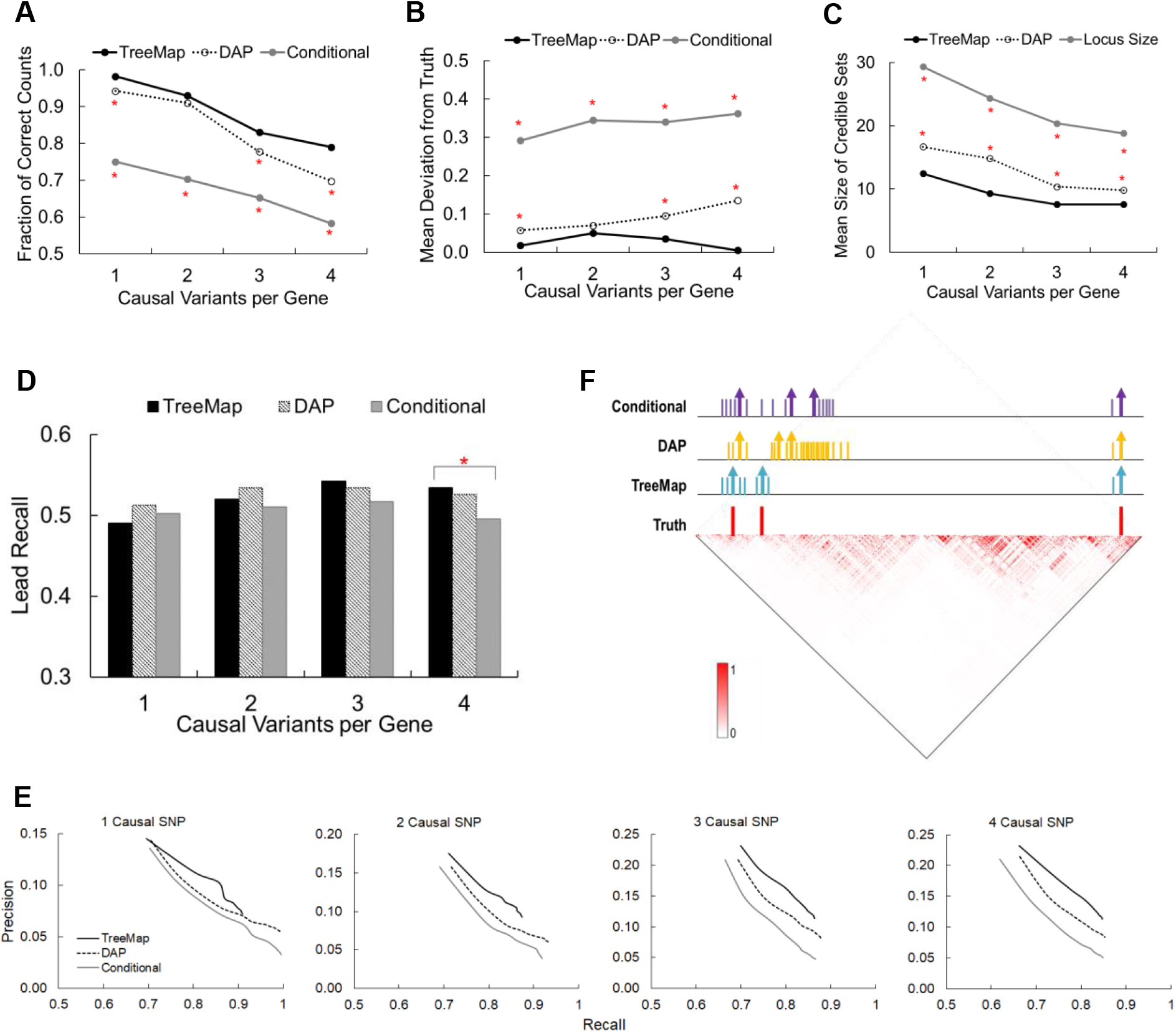
Performance of TreeMap, DAP and conditional analysis on computer simulations. (**A**) Fraction of genes with correctly predicted numbers of eQTL. Asterisks indicate that DAP or conditional analysis made significantly fewer correct predictions than TreeMap (P <0.05). (**B**) Mean deviation of the predicted and the true number of eQTL. Asterisks indicate that DAP or conditional analysis had significantly larger deviations than TreeMap (P < 0.05). (**C**) Mean size of credible sets. Because conditional analysis reports only lead variants, it is not included in the plot. Instead, average locus size of simulated causal variants is displayed that is the number of variants linked to a causal variants at r^2^>0.8. Asterisks indicate that DAP had significantly larger credible sets than TreeMap. (**D**) Recall rate among lead variants, i.e., the fraction of causal variants matched to lead variants. (**E**) Precision-recall plots for various scenarios. (**F**) An example with 3 simulated causal variants (red vertical lines) located in independent loci upstream of the *SLC28A3* gene. Vertical lines with an arrow top indicate lead eVars. Short vertical lines with a blunt top indicate linked eVars. The heat map shows pairwise r^2^ values of variants.

We then examined the sizes of credible sets (i.e., number of putative causal variants at a locus) reported by each method. A credible set contains a lead eVar and additional linked eVars. Small credible sets help narrow target candidate variants and are thus preferred. Because credible sets reported by conditional analysis contained only lead eVar, we compared TreeMap and DAP. On average, a causal variant was linked to 23 variants with r^2^≥0.8. Among these linked variants, TreeMap selected only 37%-42% to include in credible sets, whereas DAP kept 51%-61%. Therefore, the credible set of TreeMap was significantly smaller than that of DAP (all paired t tests *P*<0.05, **Fig. 2C**).

Next, we assessed how many causal variants were identified in the credible sets using two measures. The first measure is the lead recall rate (i.e., the fraction of causal variants mapped to lead eVar). In general, the lead recall rates of all three methods were similar (ranging from 49% to 54%) and varied only slightly with the number of eQTL (**Fig. 2D**). This was likely due to the relatively independence of the simulated causal variants, such that signals from multiple causal variants did not interfere with each other. However, because sampling noise could shift the signal of a true causal variant to a neighboring variant, about half of the lead eVar did not map to the causal variants. In these cases, we expected that other eVars in the credible sets should capture the causal variants. We thus assessed each method using a second measure, i.e., precision-recall curves that accounted for different sizes of credible sets. For conditional analysis, we created a credible set for each lead eVar by including linked variants with r^2^>0.8. Across all scenarios, TreeMap showed the best precision-recall curves, followed by DAP and then conditional analysis (**Fig. 2E**). To achieve a given recall rate, TreeMap had the highest precision (i.e., reporting the fewest eVars in the credible set), and conditional analysis had the lowest precision.

A representative example was simulations of 3 causal variants upstream of the *SLC28A3* gene (**Fig. 2F**). TreeMap predicted 3 eQTL correctly. At each locus, the lead eVar matched the causal variant. DAP and conditional analysis both predicted one extra eQTL and had only one lead eVar matching the causal variants. For the remaining two causal variants, conditional analysis was able to recover both from the linked variants in the credible sets, whereas DAP recovered only one and missed the other.

#### Mapping linked causal variants

We previously reported that linked causal variants concurrently regulating the transcription of the same target gene may create spurious signals on neighboring variants (**Fig. 3A**), which challenges fine-mapping (5). To simulate these cases, we generated 900 genes with two causal variants that were linked at r^2^ values >0.1 (100 genes for each r^2^ interval of 0.1 in the range of 0.1 to 1). We then examined the influence of the LD structure on the performance of each method. Overall, when the two causal variants were weakly or moderately linked (r^2^≤0.7), the impact of LD on TreeMap and DAP was mild. Both methods were able to detect two eQTL >70% of the time (**Fig. 3B**). However, when the linkage was strong (r^2^>0.7), the fraction of correct predictions quickly dropped to below 30%. When these methods made wrong predictions, they mostly collapsed the two causal variants into one eQTL (**Fig. 3C**). As expected, conditional analysis performed the worst across all scenarios.

**Figure 3.**
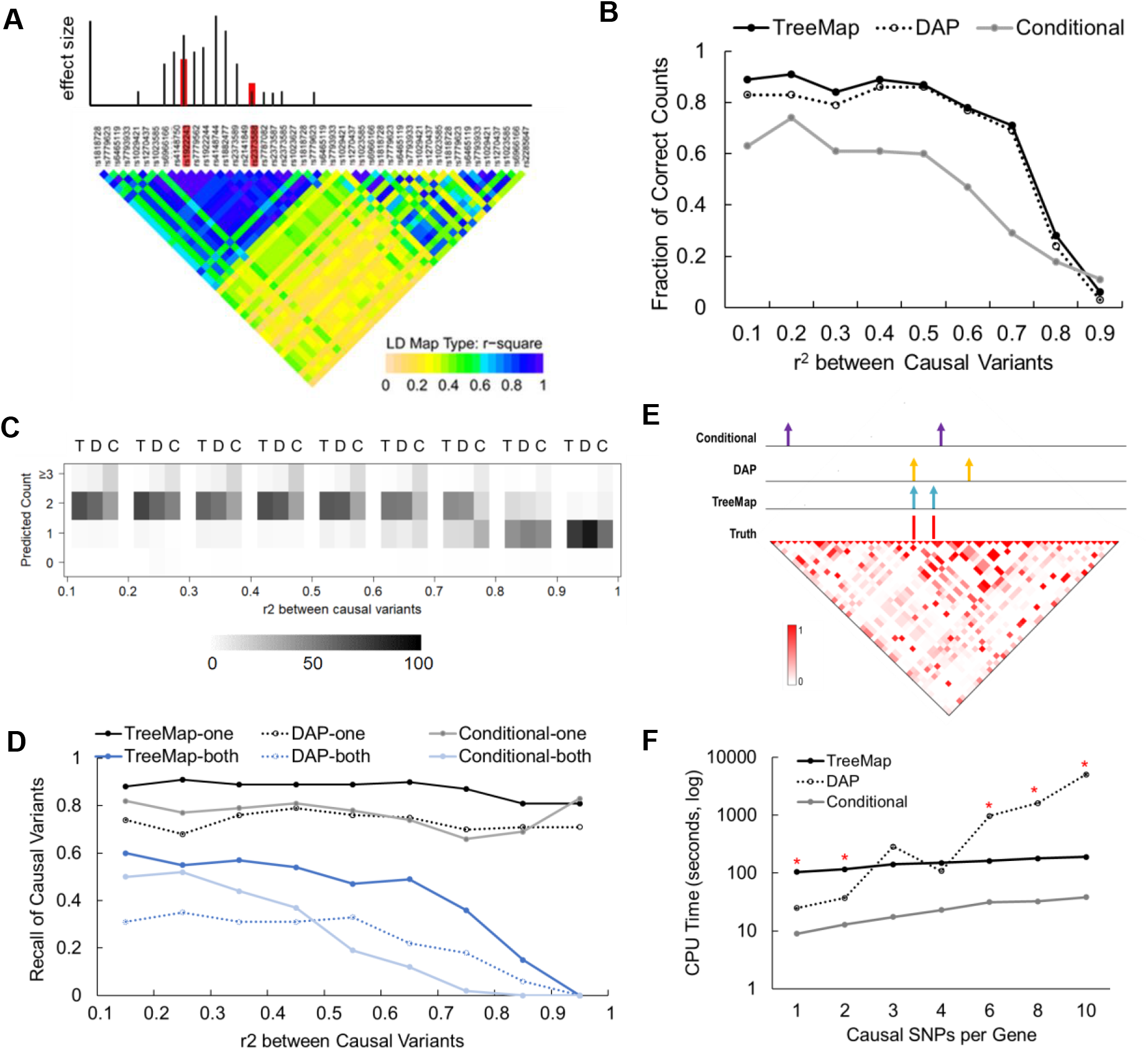
Influence of LD structure on performance. (**A**) Schematic illustration of a scenario where two regulatory variants (red bars) co-locate in an LD block, creating spurious signals (black lines) for neighboring variants. Spurious signals may be stronger than the true signals. (**B, C**) In simulated cases where two causal variants are linked, computational methods may predict two causal loci correctly, or predict fewer or more causal loci. Based on 100 simulations in each LD category, fractions of correct predictions are plotted in panel B. Numbers of predictions of zero to three causal loci are plotted in panel C. (**D**) In simulated cases where two causal variants are linked, we searched the top five eVars at each predicted locus. The rate of recalling at least one causal variant (black lines) or recalling both causal variants (blue lines) are plotted. (**E**) An example with two simulated (red) causal variants linked at r^2^=0.57 located upstream of the *SMTN* gene. Among the lead eVars (vertical lines with an arrow top) predicted by the three methods, TreeMap recalled both causal variants, whereas DAP and conditional analysis recalled 1 and 0 causal variants, respectively. Locations of lead eVars were marked. The heat map shows pairwise r^2^ values of variants. (**F**) Average CPU time (seconds in log scale) spent to analyze one simulated case.

Next, we examined if these methods could recall the two linked causal variants in a credible set. To account for different sizes of credible sets reported by each method, we limited our search among the five top-ranked eVars in each credible set. TreeMap showed the highest recall rates across the full range of r^2^ (**Fig. 3D**). As the linkage increased, the advantage of TreeMap over the other two methods became more prominent. For pairs of causal variants with r^2^ between 0.1 and 0.2, TreeMap recalled both variants in 60% of simulations, which was 10% higher than conditional analysis and 29% higher than DAP. Conditional analysis was the most sensitive to LD. Even weak to medium linkage (0.3 < r^2^ < 0.5) between the two causal variants caused the performance of conditional analysis to decline linearly. Contrarily, the performance of TreeMap and DAP were relatively stable until the LD reached a high level (r^2^ >0.7), with TreeMap showing a consistent 15% to 30% higher recall rate than DAP.

For all three methods, the recall rate of one causal variant was significantly higher than that of two causal variants (**Fig. 3D**). Again, TreeMap achieved the best recall rates across the three methods. It reported at least one causal variant among the five top-ranked eVars for 81%-91% simulations, which was on average 14% higher than the other two methods and varied only slightly across different linkage categories.

To illustrate the advantage of TreeMap, we presented a simulation in which two causal variants upstream of *SMTN* gene were linked at r^2^=0.57 (**Fig. 3E**). TreeMap correctly identified two eQTL with the lead eVars corresponding to the two causal variants. DAP also identified two eQTL. However, only one causal variant was included in its credible sets. Conditional analysis collapsed the two eQTL into a single locus and did not recall any causal variants. It also reported a false positive eQTL that was 6,496 bps away and weakly linked (r^2^=0.24) to one of the causal variant.

#### Computational efficiency

We simulated 2,000 genes, each with 1 to 10 causal variants. We distributed these causal variants randomly in the 200 kbps upstream regions of a gene. Each gene had an average of 1,700 variants genotyped in 1,835 samples. The pairwise linkages of these causal variants covered the full range of r^2^ values from 0 to 1. We then executed each method as a single-threaded process on a Dell laptop computer with an Intel^®^ Core™ i7-7600 CPU at 2.80 GHz and 16GB RAM. Conditional analysis was the most efficient method, taking an average of 9.0 seconds (secs) to analyze a gene with a single causal variant, and 38.3 secs to analyze a gene with 10 causal variants (**Fig. 3F**). To analyze a gene with only one or two causal variants, DAP took a shorter time than TreeMap (mean CPU time = 24.8 – 37.3 secs for DAP, and 104.1 – 116.3 secs for TreeMap). However, when the number of causal variants increased, the CPU time of DAP increased exponentially. For a gene with 6, 8 or 10 causal variants, DAP took an average of 965.2, 1610.6 and 5034.0 secs (16.1 minutes to 83.9 minutes) to analyze it. The CPU time of TreeMap was stable, increasing only to 162.6, 178.0 and 189.0 secs (2.7 to 3.2 minutes) in these cases.

### Applications to GTEx data

We retrieved genotype and transcriptome profiles of 123 brain hippocampus samples and 274 transverse colon samples from the GTEx data portal. There were 23,410 genes expressed in at least 10% of the brain samples and 17,065 genes expressed in at least 10% of the colon samples. For each gene, we retrieved genetic variants in a large genomic region that spanned from 2 Mbps upstream of the transcription start site (TSS) to 2 Mbps downstream of the transcription end site (TES). After removing rare variants with MAF<5%, each gene had on average 8,281 genetic variants in this region. For each gene, we applied TreeMap to organize variants into a hierarchical tree based on pairwise r^2^ values, and to identify eQTL and putative causal variants guided by the tree. To correct for multiple comparisons, we required that the primary eQTL locus of a gene had a p-value <10^−6^ corresponding to a false discovery rate of approximately 0.01 (i.e., 10^−6^ x 8,281). For auxiliary loci, we applied a lenient cutoff of p-value < 0.01 because these were post hoc tests after a significant primary eQTL was identified (23).

We detected eQTL of 4,950 genes in brain samples and eQTL of 4,636 genes in colon samples. In both tissues, only a small fraction (10%-18%) of genes had a single eQTL (**Fig. 4A**). The majority (69%-73%) had two to four eQTL. These eQTL were mostly located in noncoding regions (**Fig. 4B**, 41%-44% upstream of the target gene, 31%-38% downstream, 14%-19% intronic, 1%-2% in 5’-UTRs and 1%-2% in 3’-UTRs). Meanwhile, 3%-5% eQTL were in protein-coding regions (mean distance to TSS =5,580 bps). These distributional patterns are consistent with previous reports of eQTL from the GTEx consortium. Compared to all variants analyzed, these eQTL were >250 fold enriched in 5’-UTRs, >70 fold enriched in 3’-UTRs, >88 fold enriched in exons, and >18 fold enriched in introns (two proportions tests each having *P*<10^−8^).

**Figure 4.**
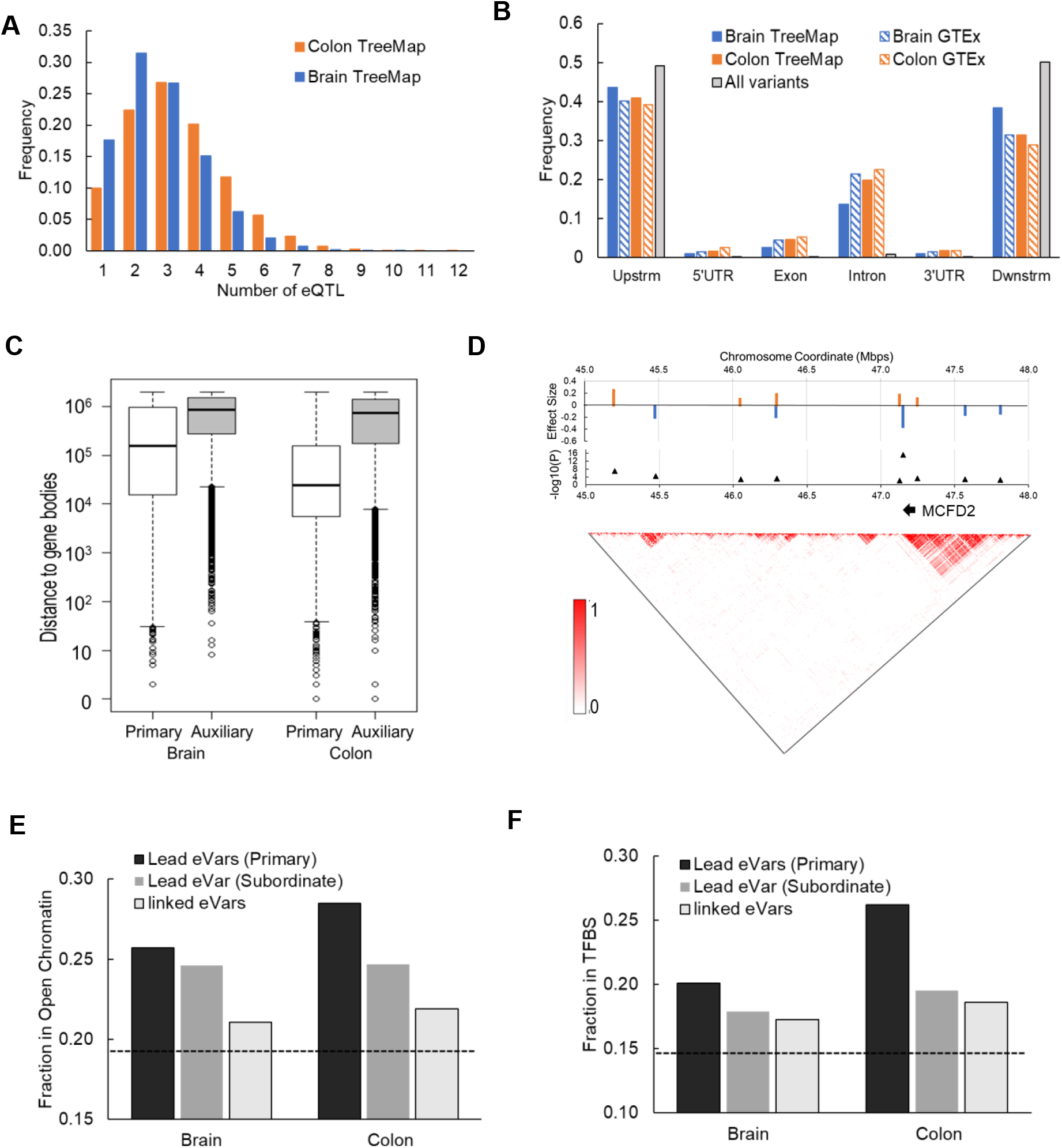
Analysis of GTEx samples. (**A**) Fractions of genes with single or multiple eQTL. (**B**) Fractions of eQTL located in various genomics regions. (**C**) Boxplots of distances to gene bodies of primary eQTL and auxiliary eQTL. (**D**) Lead eVars of eQTL of the *MCFD2* gene. Effect sizes and −log P values of each lead eVar is displayed in the top panels and LD structure of the genomic regions is displayed in the bottom panel. (**E**) Fractio of eVars in open chromatin regions. The dotted line represents the fraction of all analyzed variants located inside open chromatin. (**F**) Fractions of eVars in TFBSs. The dotted line represents the fraction of all analyzed variants located inside TFBSs.

While the identified eQTL were spread across the ±2Mbps gene-flanking regions and gene bodies, we found that primary eQTL loci were located closer to TSS or TES than auxiliary loci (median distances=150 vs. 900 kbps in brain samples, 20 vs. 74 kbps in colon samples, t-test *P*=0, **Fig. 4C**). For example, we found 10 eQTL of the *MCFD2* gene (**Fig. 4D**). The primary locus overlapped with the gene body and consisted of a lead eVar (rs34111570) and a linked eVar (rs7574514). This locus corresponded to an extensive block of LD. In fact, all eVars reported by the GTEx consortium were inside this locus. However, as we searched beyond this LD block, we found nine auxiliary loci that were located as far as 1.9 Mbps downstream of TES of this gene.

To test whether lead eVars of credible sets were more likely to be causal variants than linked variants, we examined their overlap with open chromatin regions as indicated by DNase I hypersensitivity sites, and overlap with transcription factor binding sites (TFBSs) as annotated in the ENCODE database. As expected, the fraction of variants in open chromatin regions was highest among lead eVars in primary eQTL (26%-29) and lower in linked eVars (21%-22%, two proportions test *P*<10^−15^, **Fig. 4E**). Furthermore, all eVars are enriched in open chromatin regions as compared to all variants analyzed (19%, hypergeometric test *P*<10^−26^). Similarly, the fraction of variants in transcription factor binding-sites (TFBSs) was the highest among lead eVars at primary eQTL (18%-26%) and lower among linked eVars (18%-19%, two proportions test *P*<10^−8^), both of which were significantly higher than that among all variants analyzed (15%, *P*<10^−20^, **Fig. 4F**).

We found shared eQTL for 1,377 genes in both brain and colon samples, 739 (53.7%) of which had the same putative causal variants. When these putative causal variants did not overlap, most of them (397 among 638) were in the same LD block (**Fig. 5A**) or in close vicinity (**Fig. 5B**). TreeMap identified many eVars located far from gene bodies not explored by other methods. This was expected because TreeMap searched up to 4Mbps regions for eQTL in regions that had generally not been analyzed before. However, if these distal eVars were found in both tissues and shared close genomic positions or LD blocks, they were more likely to be functional. For example, the primary eQTL of the *AC018804* gene in brain and colon samples were located 1.3 Mbps downstream of the gene. The lead eVars had extraordinary p-values (10^−17^ and 10^−28^) and concordant effect sizes (0.79 and 1.27) in brain and colon samples, respectively. The two lead eVars were within a 7.2 kbps interval on chromosome 3 (132,240,509 in brain samples and 132,233,317 in colon samples). Furthermore, both lead eVars were in open chromatin regions, providing additional evidences of their functional roles.

**Figure 5.**
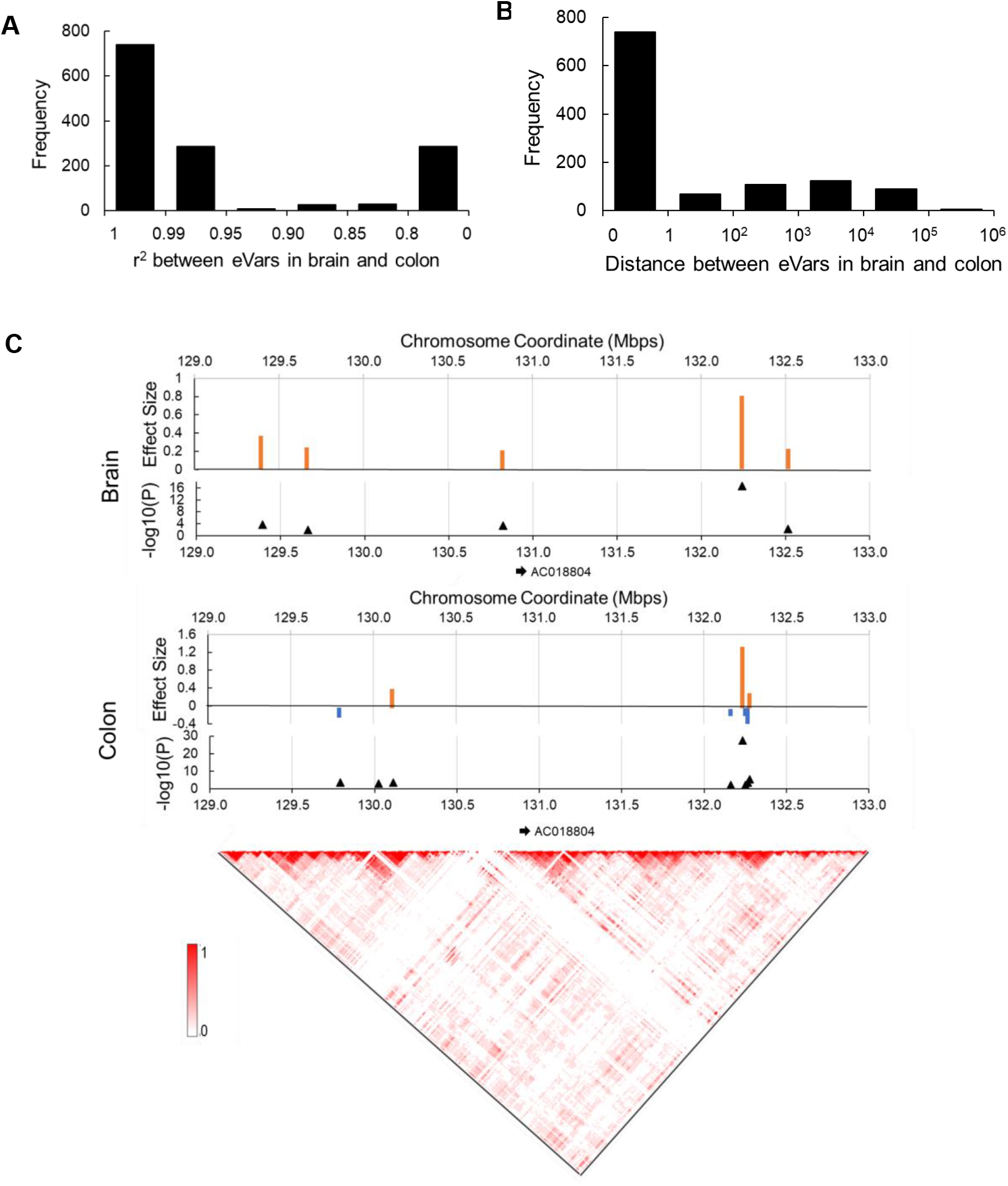
Overlapping eQTL between brain and colon samples. (**A**) Numbers of genes sharing eVars in the same LD blocks. LD blocks were defined based on r^2^ values. (**B**) Numbers of genes sharing nearby eVars. (**C**) eQTL of the *AC018804* gene in brain and colon samples. The primary eQTL in both tissues is located 1.3 Mbps downstream of the gene. The lead eVars were within 7.2 Kbps of one another on chromosome 2 (132,240,509 in brain samples and 132,233,317).

## DISCUSSION

With increasing sample sizes for eQTL mapping it has become apparent that the majority of all genes have a complex pattern of regulation influenced in cis by multiple SNPs. Fine mapping of the causal variants is constrained by the high degree of LD covering most regulatory regions, and high levels of polymorphism such that credible intervals average 100 sites or more (4). Three broad approaches to dealing with this complexity are being developed: stepwise conditional regression, Bayesian dimensionality reduction, and haplotype-based modeling. The method introduced in this study, TreeMap, combines elements of the latter two.

An important aspect of haplotype-based methods is the heuristic definition of haplotypes. Perhaps the most rigorous procedure uses the four-gamete test to identify minimal length haplotypes by virtue of inferred recombination events. A genome-wide association mapping method based on this approach, HaploSNP (24), explains much more of the variance per locus. However, because haplotypes are greatly susceptible to biases introduced for example by population structure and do not have base pair resolutions, HaploSNP has not been widely adopted for fine mapping. Instead, we here propose a hierarchical approach based on successive LD thresholds. Causal variants are assumed to be embedded in LD blocks although the extents of linkage are unknown. Transcriptional effects are gleaned from comparison of the likelihoods of models for blocks defined by varying LD thresholds. This algorithm is thus independent of cladistic methods for assembly of cladograms with ad hoc thresholds that may have hampered adoption of earlier iterations of haplotype-based approaches (17,24).

Using extensive simulation, we show that TreeMap modestly, yet significantly, outperforms representative alternative multi-site eQTL mapping algorithms (stepwise conditional regression, and DAP) in several key regards. First, it recovers more independent variants, particularly as the complexity of multisite regulation increases. Second, it reduces the size of the credible interval as assessed by improvement in the precision-recall curve. Third, it recovers more causal variants under LD. Furthermore, since the method is computationally far less demanding than even the fastest Bayesian approach, DAP, it is possible to scan 4Mb around the transcription start site, and this doubling of the potential regulatory region led to the discovery of multiple hitherto unrecognized distal eQTL in the GTEx dataset.

There remain several limitations to be addressed. Like the other methods, performance drops as the number of independent causal variants in an eQTL increases, particularly if they fall within intervals of high LD. Under soft selection scenarios, it may be expected that regulatory regions will harbor more than one variant influencing gene expression, with multiple signals embedded in a haplotype. Variants that have opposing signs of effect will tend to reduce the overall signal. Methods for multi-site mapping of tightly linked causal variants need to be further explored. One strategy is to incorporate functional evidence from ENCODE or evolutionary conservation into the mapping algorithm (25,26), though this strategy seems to be more productive for promoter-proximal than distal elements.

TreeMap, and future improvements of incorporating prior biological knowledge in fine mapping algorithms, facilitate discoveries of regulatory variants from genome-association studies and whole-genome sequencing studies as well. The capability of searching long genomic regions makes it a promising approach to identifying novel distal regulatory variants underlying human diseases and other health-related genotypes.

## Data availability

GTEx data are accessible from the GTEx portal (https://gtexportal.org). All simulation data used in this study are available at the TreeMap Github site (https://github.com/liliulab/treemap).

## ACKNOWLEDGEMENT

We thank Dr. Panwen Wang for insightful discussions.

